## Supplemental Text for "When to wake up? The optimal waking-up strategies for starvation-induced persistence"

#### Contents

**1 Relaxing the assumption 3**
1.1 Non-zero killing rate at the full dormant state . . . . . . . . . . . 3
1.2 A gradual growth resurrection model . . . . . . . . . . . . . . . . 4

**2 The threshold of the extinction 5**

**3 The equation used for the evolution simulation of the lag time 7**

**4 A multi-step model 9**

**5 The computational procedure for obtaining the optimal combined-Erlang distributions 10**

**6 Mapping to the Kussell-Leibler model 11**

**7 The critical $\gamma$ and $p$ 13**

**8 A condition for a discontinuous transition for arbitrary probability distribution functions 16**

**9 Optimal lag time distributions 18**
10.3 A sufficient and necessary condition for the discontinuous transition 32

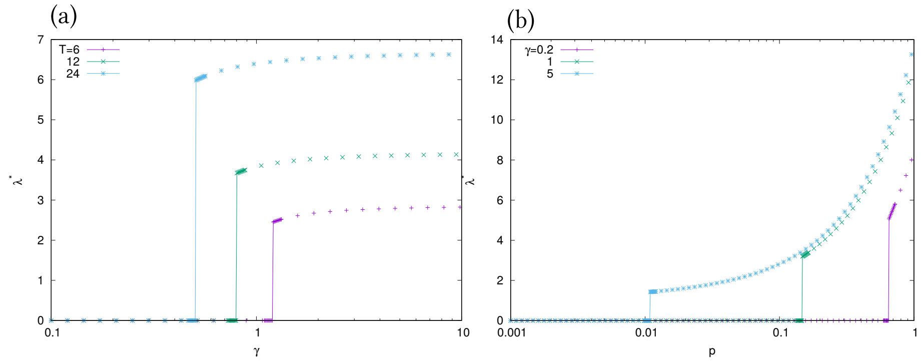

Fig.S1: **The effect of changing** $\gamma$ (a). The transition triggered by $\gamma$. (b). The transition triggered by $p$ with different $\gamma$ values. $p = 0.2$ for (a) and $T = 12$ for (b).

Note: For the consistency with the results shown in the main text, throughout the present document we define the Dirac's delta function peaked at the origin $\delta(x)$ so that

$$\int_0^\infty \delta(x)dx = 1$$

is satisfied.

### 1 Relaxing the assumption

In this section, we relax the assumption that the growth and death take place only in the single, growing state. We first specifically see what happens if the cells in the dormant state are killed by the antibiotics. Next, we study the model where the cells recover their growth and death rate gradually.

#### 1.1 Non-zero killing rate at the full dormant state

Here we introduce the non-zero killing rate also for the full dormant state. We assume that the killing rate at the full dormant state is smaller than that of the active state, and thus, we set it to $\alpha\gamma$ where $0 < \alpha < 1$. The rate equation is given as

$$\dot{d}(t) = \begin{cases} -(\alpha\gamma + 1/\lambda)d(t) & (t < T) \\ -d(t)/\lambda & (t > T) \end{cases} \quad (1)$$

$$\dot{g}(t) = \begin{cases} d(t)/\lambda - \gamma g(t) & (t < T) \\ d(t)/\lambda + g(t) & (t > T) \end{cases} \quad (2)$$

The fitness function are given by

$$\begin{aligned} \tilde{F}_I(\lambda, \alpha, \gamma, p, T) &= (1-p)\ln\left[\frac{1}{1+\lambda}\right] \\ &+ p\left(-T + \ln\left[\frac{1}{1-(1-\alpha)\gamma\lambda}\left(e^{-\gamma T} - e^{-(1/\lambda+\alpha\gamma)T}\right) + \frac{1}{1+\lambda}e^{-(1/\lambda+\alpha\gamma)T}\right]\right). \end{aligned}$$

Fig. S2(a) shows the optimal lag time as the function of $\alpha$ and $T$. The optimal $\lambda$ value still shows the transition by changing $T$. Since the nature of the transition is the same with $\alpha = 0$ case, the transition is triggered by changing the severeness of the antibiotics application by changing either $p$, $\gamma$, and $T$. Also the model shows transition by changing $\alpha$ in the region $T \gtrsim 10$. The transition takes place discontinuously as shown in Fig. S2(b). Note that the optimal lag time for $\alpha = 1$ case is zero regardless of the other parameter values because there is no reason to stay at the dormant state.

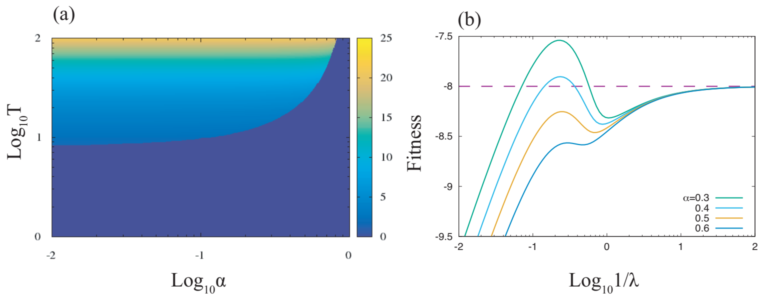

Fig.S2: **The optimal lag time for non-zero killing rate at the full dormant state** (a). The optimal $\lambda$ is plotted as a function of $\alpha$ and $T$. $\lambda^*$ shows the discontinuous transition. (b). The fitness function for several choices of $\alpha$. $\lambda^*$ transits in the same way as Fig1(b) in the main text. The dashed line represents $\tilde{F}_I(\lambda = 0, \alpha, \gamma, p, T)$. $p = 0.2$, $\gamma = 1.0$, and $T = 20$ for (b).

#### 1.2 A gradual growth resurrection model

We relax the assumption that cells recover the ability for growth together with death abruptly when they reached to the single growing state. To making a gradual recovery of the growth and death rate possible, we use the model with multiple states.

Here we consider the model has $M + 1$ states ($M$ steps). Each state has the growth, death, and transition rate to next state, $\mu_i, \gamma_i$, and $1/\lambda_i$. Since the scope of this section is an impact of the gradual recovery of the growth and the death rate on the main result, here we assume that $\lambda_i$'s are uniform among all the states while it is zero at the final, the $M$th state. Then, the temporal evolution of the population after an inoculation is ruled by

$$\frac{d}{dt}N_i(t) = \left(\hat{\delta}_{i,0}N_{i-1} - \hat{\delta}_{i,M}N_i\right)M/\lambda + \alpha_i(t)N_i, \ (0 \leq i \leq M)$$

$$\alpha_i(t) = \begin{cases} -\gamma_i & (t < T) \\ \mu_i & (t \geq T) \end{cases}$$

where $N_i$ represents the population of the cells at the $i$th state and $\hat{\delta}_{i,j}$ is the complementary of the Kronecker's delta defined as $1 - \delta_{i,j}$. We set $\mu_M$ as unity. The solution of the ordinary differential equation is given by

$$N_i(t) = \sum_{j=0}^{i} \xi_{ij} c_j e^{\beta_j t M/\lambda} \quad (3)$$

where, parameters are given by

$$\xi_{ij} = \begin{cases} \prod_{k=j+1}^{i} (\beta_j - \beta_k)^{-1}, & (i > j), \\ 1 & (i = j) \end{cases}$$

$$\beta_j = \alpha_j M \lambda - \hat{\delta}_{j,M},$$

and $\xi_{ij}'$s with $i < j$ do not appear in Eq.(3). We put a superscript $\pm$ to $N_i, \xi_{ij}, \alpha_i, \beta_i$, and $c_i$ representing before($-$) and after ($+$) finishing the antibiotics application. $c_i^{\pm}$'s are determined to satisfy the initial condition $N_0^-(0) = 1, N_1^-(0) = \cdots = N_M^-(0) = 0$ and continuity of the solution at the end of the antibiotics application, $N_i^-(T) = N_i^+(T)$.

The antibiotics-free solution is obtained by setting $T = 0$. Since the definition of the fitness of a single round is the logarithmic growth of the population under a large $t$ limit, relative to the zero- antibiotics application time and lag time, it is given as

$$f(T) = \lim_{t \to \infty} \frac{\sum_{i=0}^{M} \sum_{j=0}^{i} \xi_{ij}^+ c_j^+ e^{\beta_j^+ tM/\lambda}}{e^t} = c_M^+,$$

and accordingly, the fitness function is calculated. The fitness function is plotted against $\lambda$ for several choices of $p$ values and $M = 2$ and 4 in Fig. S3. The fitness function has its maximum at $\lambda = 0$ when $p$ is small, while it forms local maximal which exceeds the fitness value at the origin as $p$ increases. The result suggests the robustness of the discontinuous transition described in the main text against the model extension that the cells recover their growth rate and antibiotic susceptibility.

### 2 The threshold of the extinction

Here we study the slightly modified model in which the population cannot be lower than a threshold value. Note that the populations ($d(t)$ and $g(t)$ in Eq.(1) and (2) in main text) decrease only in the duration of the antibiotics application, and the shorter the lag time $\lambda$ is, the more prominent the killing effect gets. If $\lambda$ is sufficiently larger than $T$, the total population $d(t) + g(t)$ never reaches to the threshold. Therefore, the introduction of the threshold sets the lower limit of the lag time $\lambda$ so that all the cells are not killed by the antibiotics.

To guarantee the population is greater than a threshold denoted by $\delta_{\text{ext}}$, the inequality

$$d(T) + g(T) = \frac{\lambda\gamma}{\lambda\gamma - 1} e^{-T/\lambda} - \frac{1}{\lambda\gamma - 1} e^{-\gamma T} > \delta_{\text{ext}} \tag{4}$$

needs to be satisfied. We set $\delta_{\text{ext}} = 10^{-6}$ because the initial value of $(d + g)$ is normalized to unity in the main text and typical numbers of the bacteria in a test tube for a few milliliters range from $10^6$ to $10^9$. By substituting $\lambda = 0$ into Eq.(4), we get the condition for allowing zero lag time, given as $e^{-\gamma T} > \delta_{\text{ext}}$. If

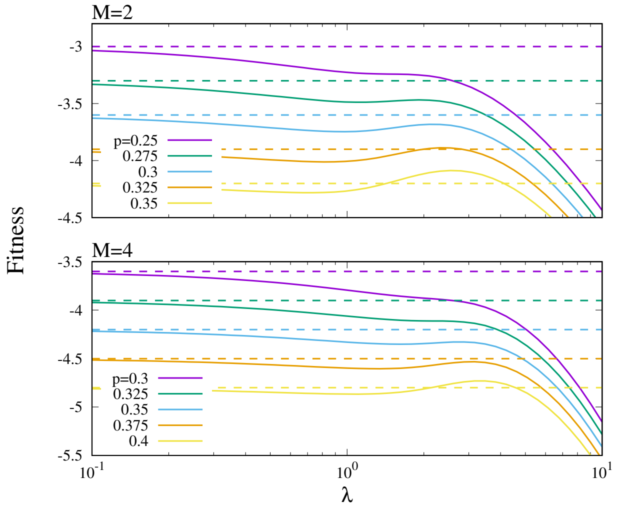

Fig.S3: **The fitness values of the gradual resurrection model** The fitness values of the $M$ steps, gradual resurrection model are plotted against $\lambda$ for several values of $p$ where $M = 2$ (top) and 4 (bottom). The dashed lines represents the fitness value at $\lambda = 0$ of the corresponding color. As $p$ value increases, the fitness function forms a peak and the value at the peak exceeds the value at $\lambda = 0$. Here $\mu_i$ and $\gamma_i$ are given as $\mu_i = \gamma_i = i/M,\ (0 \leq n \leq M)$, and $T = 6.0$.

this condition is not satisfied, the minimum lag time $\lambda_{\text{min}}$ is set by Eq.(4) with the replacement of inequality with equality.

Fig. S4 shows $\max\{\lambda^*, \lambda_{\text{min}}\}$ in $(T, p)$ plane where $\lambda^*$ denotes the optimal average lag time without the restriction (Eq.(4)). The plane is divided into four regions: (i). the zero lag time is optimal and feasible, (ii). the zero lag time is optimal but $\lambda_{\text{min}}$ is non-zero, (iii). $\lambda_{\text{min}}$ is non-zero but the optimal lag time is greater than that, and (iv). the optimal lag time is non-zero and $\lambda_{\text{min}} = 0$. The white line gives the boundary between $\lambda^* = 0$ and $\lambda^* > 0$, and the pink line being given as $-\ln(\delta_{\text{ext}})/\gamma$ divides the region into $\lambda_{\text{min}} = 0$ and $\lambda_{\text{min}} > 0$ part.

The figure tells that, by changing $p$ as the parameter, one can observe the discontinuous transition for any reasonable $T$ value while the jump size of the optimal lag time gets less as $T$ becomes larger. On the other hand, when $T$ is chosen as the parameter, one can observe only the continuous transition in a reasonable timescale with a rare antibiotics application ($p$ as less than a few percent).

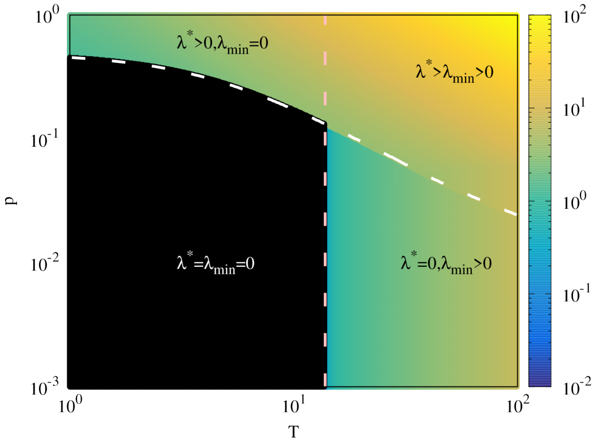

Fig.S4: **The optimal lag time** The heatmap of the optimal lag time. The white and pink dashed lines divide the space into four regions. Color shows $\max\{\lambda^*, \lambda_{\text{min}}\}$. $\gamma$ is set to unity.

### 3 The equation used for the evolution simulation of the lag time

The system of equations used for the evolution simulation (in the section III-C) is following

$$
\begin{aligned}
\frac{d}{dt}d^{(i)}(t) &= -d^{(i)}(t)/\lambda_i \ (0 \le i < N) \\
\frac{d}{dt}g^{(0)}(t) &= d^{(0)}(t)/\lambda_0 + (1-\epsilon)\alpha g^{(0)}(t) + \epsilon\alpha g^{(1)}(t) \\
\frac{d}{dt}g^{(i)}(t) &= d^{(i)}(t)/\lambda_i + (1-2\epsilon)\alpha g^{(i)}(t) + \epsilon\alpha(g^{(i-1)}(t) + g^{(i+1)}(t)), \ (1 \le i \le N-2) \\
\frac{d}{dt}g^{(N-1)}(t) &= d^{(N-1)}(t)/\lambda_{N-1} + (1-\epsilon)\alpha g^{(N-1)}(t) + \epsilon\alpha g^{(N-2)}(t),
\end{aligned}
$$

where $d^{(i)}$ and $g^{(i)}$ represents the population of the $i$th-type cells in the dormant, and the growing state, respectively. $(\alpha, \epsilon)$ is $(-\gamma, 0)$ if the antibiotics is applied and $t < T$, otherwise, it is $(1.0, 10^{-3})$. $\lambda_i$ is given as $i\Delta\lambda + 10^{-6}$.

The effect of the value of $\Delta\lambda$ is shown in Fig. S5. For the sharp transition of the average lag time, the choice of $\Delta\lambda$ is not crucial as long as it is reasonably small. When $\Delta\lambda = 0.5$ is chosen, the lag time changes rather continuously due to the large fluctuations.

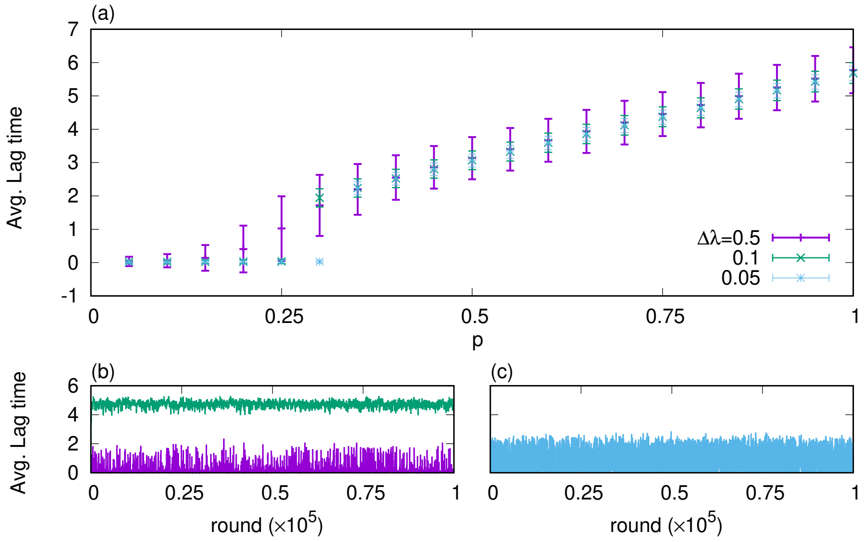

Fig.S5: **The effect of** $\Delta\lambda$ (a). Time average of population-averaged lag time for three choices of $\Delta\lambda$. (b) and (c). The population-averaged lag time at each round with $\Delta\lambda = 0.5$. $p = 0.15$ and $0.8$ for (b) and $0.4$ for (c). The same parameter values with Fig.1(c) and (d) in the main manuscript are used for the rest.

Also, we changed the rule of the mutation so that any single population can mutate to any other population because some mutations may change the lag time drastically. Equation is then given as

$$\begin{aligned} \frac{d}{dt}d^{(i)}(t) &= -d^{(i)}(t)/\lambda_i \ (0 \le i < N) \quad (5) \\ \frac{d}{dt}g^{(i)}(t) &= d^{(i)}(t)/\lambda_i + (1-\tilde{\epsilon})\alpha g^{(i)}(t) + \sum_{j \ne i} \frac{\tilde{\epsilon}}{N-1}\alpha g^{(j)}(t), \ (0 \le i < N). \end{aligned}$$

As the noise level by the mutation has been increased dramatically, dynamics become much noisier than the previous model, and thus, the averaged lag time ($\lambda_{\text{avg.}}$) shows rather continuous change with $p$ (Fig. S6(a)). However, looking at the population-averaged lag time at each round, it stays either the lower side ($\lambda_{\text{avg. single round}} \approx 10^{-2}$) or the higher side ($\lambda_{\text{avg. single round}} \approx 10$) most of the time as shown in Fig. S6(b). Thus, the continuous behavior stems from the continuous change of the residual time of the two branches rather than that the stable lag time changes continuously with $p$.

### 4 A multi-step model

As an extension of the model (Eq.(1) and (2) in main text), we consider the model with multiple dormant states. All the cells start to wake up from the 0th dormant state after the re-inoculation, and transit one by one through the $M$ dormant states to regain the growth rate at the growing state. Here, we assume that the growth rate and the death rate are non-zero only in the growing state, and the transition rates from one state to the next state are uniform (this is a special case of the gradual resurrection model). Then, the temporal evolution of the population after an inoculation is ruled by

$$\frac{d}{dt}d_0(t) = -d_0(t)M/\lambda, \quad (6)$$

$$\frac{d}{dt}d_i(t) = (d_{i-1} - d_i(t))M/\lambda, \ (1 \le i \le M-1) \quad (7)$$

$$\frac{d}{dt}g(t) = \begin{cases} d_{M-1}(t)M/\lambda - \gamma g(t) & (t < T) \\ d_{M-1}(t)M/\lambda + g(t) & (t > T), \end{cases} \quad (8)$$

where $M/\lambda$ is the rate of the transition from the $i$th to the $i+1$th state.

This dynamics leads to Erlang distribution $P(l) = l^{M-1}e^{-l/\lambda_M}/((M-1)!\lambda_M^M)$ as the lag time distribution where $\lambda_M = \lambda/M$. The average lag time is $\lambda$. The single-round fitness $f_M(T)$ is now given by

$$\begin{aligned} f_M(T) &= e^{-(1+\gamma)T}(1-\gamma\lambda_M)^{-(1+M)} \quad (9) \\ &+ e^{-(1+1/\lambda_M)T} \sum_{n=1}^{M+1} \frac{(T/\lambda_M)^{M+1-n}}{(M+1-n)!}\Big((1+\lambda_M)^{-n} - (1-\gamma\lambda_M)^{-n}\Big). \end{aligned}$$

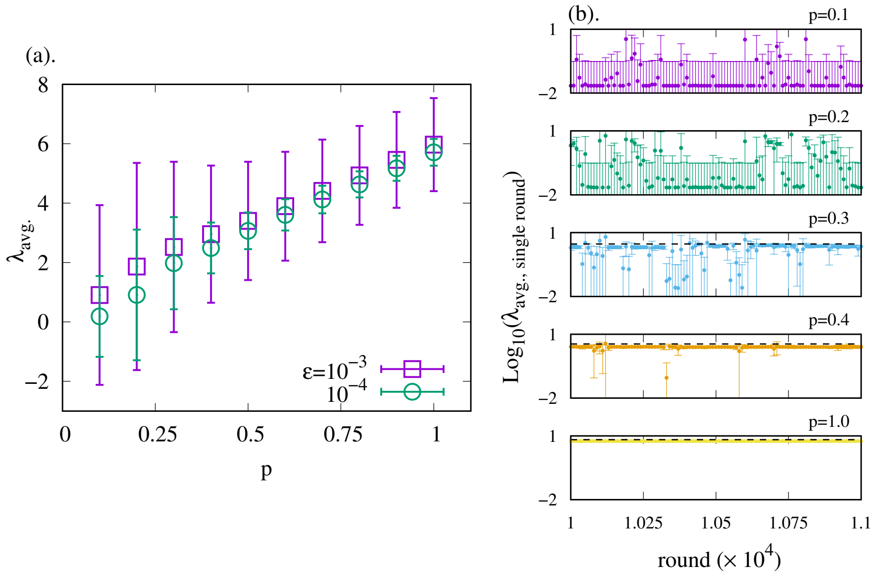

Fig.S6: **Random mutation model** (a). Time average of population-averaged lag time of the random mutation model (Eq.(5)) for two choices of $\tilde{\epsilon}$. (b) The population-averaged lag time with one standard deviation of each round (from $10^4$th to $1.1 \times 10^4$th round) for several $p$ values. Data points are plotted every 10 rounds. Error bar indicates the standard deviation. The black dashed line representing the optimal lag time obtained from the optimization of $F_I$ is added for each panel if the optimal lag time is non-zero ($p = 0.3, 0.4$ and $1.0$). $N = 200, \Delta\lambda = 0.1$,$T = 6.0$ and $\gamma = 1.0$. $\tilde{\epsilon} = 10^{-4}$ for (b).

As described in S1 text section 2, $M$-step models, including the main model ($M = 1$ case) in the manuscript, generally exhibit the discontinuous transition of $\lambda$ as long as the lower bound of the support of $q(T)$, a distribution of the antibiotics application time, is non-zero. It is worth mentioning that as $M$ increases, the optimal fitness value becomes greater if $T$ is fixed because the Erlang distribution with a large $M$ is peakier. However, if $T$ is distributed, a large $M$ is not always beneficial, and there typically optimal $M$ exists.

### 5 The computational procedure for obtaining the optimal combined-Erlang distributions

For obtaining the optimal lag time distribution of the sequential model with $N_p$ phenotypes and $M$ steps (Fig.6 in main text), we carried out following computations.

For given $N_p$, $M$, and other parameters $(p, \gamma$, and $q(T))$, the fitness value

$$F = \int_0^{\mathrm{Tmax}} \hat{q}(T) \ln\Big[\sum_{i=0}^{N_p-1} x_i f_M(T;\lambda_i)\Big] dT,$$

is maximized where $\hat{q}(T) = (1-p)\delta(T) + pq(T)$. $f_M(T;\lambda_i)$ is given in Eq.(9) while $\lambda$ is replaced by $\lambda_i$, and $x_i$ represents the fraction of the $i$th phenotype. Here, the variables for the optimization are $x_i'$s and $\lambda_i'$s and the restriction $\sum_{i=0}^{N_p-1} x_i = 1$ is introduced.

### 6 Mapping to the Kussell-Leibler model

The focus of the paper written by Kussell and Leibler [1] is to ask the optimal strategy of the bacterial population which has multiple types (either genotypic or phenotypic) to grow most efficiently under the fluctuating environment. The interactions among cells are not considered in the model, and thus, the model is given as the linear ordinary differential equations, $\dot{\mathbf{x}}(t) = A_k \mathbf{x}(t)$ where $\mathbf{x}$ represents the population vector at time $t$. The matrix $A_k$ consists of the growth term, and in addition, the transition term among the types representing either the genetic mutation or phenotypic switch in the environment $k$. The matrix changes from $k$ to $k'$ $(k \neq k')$ after a certain period of time has passed to emulate the fluctuations of the environment. The growth rate of the $i$th species (whose population is represented by the $i$th column of $\mathbf{x}$) has a different value depending on the environment index $k$. It can be either positive or negative. The negative growth rate then indicates that the $i$th type cannot even survive in the environment $k$ and dies over time. Because of the environmental dependency of the growth rate, the population needs to tune the transition rate among the types so that the total number of the cells increases well.

The authors studied two scenarios, namely the responsible adaptation and the stochastic adaptation. In the responsible adaptation, the cells can "sense" the environment and use the environment-specific transition rates. In contrast, in the stochastic adaptation, the cells are allowed to use only a set of transition rates independently of the environment. One of the main results of the paper is that the optimal strategy for stochastic adaptation is mimicking the fluctuation of the environment. For instance, if the number of the types and the number of environments are the same and the $i$th type has the highest growth rate in the $i$th environment, the optimal transition rate of the $i$th to the $j$th species, $H(i \to j)$ is given as $b(i \to j)/\tau_i$ where $b(i \to j)$ is the transition rate from the $i$th to $j$th environment, and $\tau_i$ represents the latency time of the $i$th environment. The optimal lag time distribution is not the complete copy of the distribution of the antibiotics application time, and thus, there should be mathematical differences between the KL model and our model.

To show how to map our model to this framework and where the different consequence comes from, we briefly explain the analysis done by the authors. Since the solution of the ordinary differential equation is given as $\mathbf{x}(t) = \exp[A_k t]\mathbf{x}(0)$, we obtain a sequence

$$\begin{aligned} \mathbf{x}_{\mathrm{ini}} &\to e^{A_{\mathcal{E}_0}\tau}\mathbf{x}_{\mathrm{ini}} \to e^{A_{\mathcal{E}_1}\tau}e^{A_{\mathcal{E}_0}\tau}\mathbf{x}_{\mathrm{ini}} \to \cdots \\ &\to \prod_{i=0}^{N} e^{A_{\mathcal{E}_i}\tau}\mathbf{x}_{\mathrm{ini}} \to \cdots \end{aligned}$$

representing the total population just before the $i$th change of the environmental condition, where $\mathcal{E}_i$ represents the environment at the $i$th round. Here we eliminated the environment dependency of the latency time $\tau$ because the dependency is unnecessary for the later arguments as long as $\tau$ is sufficiently large.

The analysis in the paper is calculating the average Lyapunov exponent over the sequential environmental changes and maximizing it by modulating the transition rate among the types. Thus, by denoting the population vector after the $n$th round as $\mathbf{x}_n$, the question is formulated as the optimization problem of

$$F_{KL} = \left\langle \frac{\mathbf{1}\cdot e^{A_{\mathcal{E}_{n+1}}\tau}\mathbf{x}_n}{\mathbf{1}\cdot\mathbf{x}_n} \right\rangle \tag{10}$$

where the average $\langle\cdot\rangle$ is taken over $n$'s and $\mathbf{1}$ is the vector having the same dimension with $\mathbf{x}_n$ given as $\mathbf{1} = {}^t(1, 1, \cdots, 1)$. The optimal transition rates among the types are calculated by using the perturbation method. The population growth in a single round is determined only from the growth rates, while the transition rates play a role in modulating the population distribution among the types. The central assumption is that there is a separation among the eigenvalues of $A_k$ to simplify the evaluation of the fitness in a single round. If the assumption holds, the fitness is well-approximated by the highest growth rate in the environment $k$ and the population of the fittest type under the large $\tau$ limit. When the assumption is violated, the contribution of all the other types are unignorable, and thus, the simple argument is no longer valid.

Now, we describe how to map our model to this framework. We consider the generic case that the population of the cells has a lag time distribution $r(l)$. We set the upper limit of the distribution $T_{\max}$ and discretize the distribution by a certain size of the bin to make a finite number of sectiongroups having the lag time $l_i$. Now, the different "types" in the KL framework corresponds to the groups of the cells with different lag time and the different environment means the different antibiotics application time $T_k$. We suppose that the number of types and the number of the environment are equal for simplicity.

Our population dynamics is non-autonomous because the cells of the $i$th type are in quiescent until $t = l_i$. However, as long as $T_{\max} < \tau$ holds, we can write down the autonomous, effective population dynamics given as

$$\dot{\mathbf{y}}(t) = B_k\mathbf{y}(t) \tag{11}$$

with

$$B_k = E - \mathrm{diag}\Big(\ln[f(l_0, T_k)]/\tau, \cdots, \ln[f(l_{N-1}, T_k)]/\tau\Big)$$

where $E$ is the unit matrix and $\mathbf{y}(t)$ is the population vector at time $t$ and $\ln f(l_i, T_k)$ represents the population loss by having the lag time $l_i$ and exposed to the antibiotics for $T_k$. Note that the number of the $i$th type cells at time $\tau$ is given as $f(l_i, T_k)\exp[\tau]y_i(0)$ being consistent with the original population dynamics in the main text. Now, we can emulate our population dynamics in the KL framework by choosing the discretized lag time distribution $\mathbf{r}$ as the initial vector $\mathbf{y}(0)$. Note that Eq.(10) is defined for an arbitrary population vector $\mathbf{x}_n$, and thus, starting from the same vector $\mathbf{r}$ every time is a special case of Eq.(10). Also, Eq.(10) is invariant for the normalization of the total population at every round.

Now, we mapped the present model to the KL framework, and thus, are able to see where the difference between the prediction in ref. [1] and the present result comes from. The critical difference is that the separation of the eigenvalue never happens under $\tau \to \infty$ limit in the present model. The growth rate correspondence in the present model, $1 - f(l_i, T_k)/\tau$ lose the $i$-dependence under this limit. Thus, there is no way to evaluate the population growth by only a single eigenvector, and it leads to a different consequence.

### 7 The critical $\gamma$ and $p$

In this section, we describe detailed calculations. For the sake of readability, we first list the notations of some of the functions and parameters.

$$
\begin{aligned}
F_I(\lambda, \gamma, p, T) &= p\Big(-T + \ln\Big[\frac{1}{\gamma\lambda - 1}\Big(e^{-T/\lambda} - e^{-\gamma T}\Big) + \frac{1}{1+\lambda}e^{-T/\lambda}\Big]\Big) + (1-p)\ln\Big[\frac{1}{1+\lambda}\Big]. \\
F_I^{\infty}(\lambda, p, T) &\equiv \lim_{\gamma\to\infty} F_I(\lambda, \gamma, p, T) = -pT(1+\lambda^{-1}) - \ln[1+\lambda]. \\
\lambda^*(\gamma, p, T) &\equiv \arg\max_{\lambda} F_I(\lambda, \gamma, p, T). \\
\lambda^*_{\infty}(p, T) &\equiv \arg\max_{\lambda} F_I^{\infty}(\lambda, p, T) = 2\left(-1 + \sqrt{1 + \frac{4}{pT}}\right)^{-1}
\end{aligned}
$$

First we show there is a critical $0 < p < 1$. The first-order derivative of the fitness function respect to $\lambda$ is given as

$$
\frac{(1+\lambda)}{p}\frac{\partial F_I}{\partial \lambda} = -\frac{1-p}{p} + (1+\lambda)\frac{\partial}{\partial \lambda}\ln\left[\frac{1}{\gamma\lambda - 1}\Big(e^{-T/\lambda} - e^{-\gamma T}\Big) + \frac{1}{1+\lambda}e^{-T/\lambda}\right].
$$

As $p \to 0$, the first term of the right hand side of the equation diverges to $-\infty$ while the second term is constant, the optimal $\lambda$ is zero. Therefore, the transition of the optimal $\lambda$ triggered by $p$ happens when $(1-p)/p$ approaches to the second term from above to make an intersection. This intersection always exists as long as $\gamma > 0$ because

$$
\lim_{\lambda\to+0}(1+\lambda)\frac{\partial}{\partial \lambda}\ln\left[\frac{1}{\gamma\lambda - 1}\Big(e^{-T/\lambda} - e^{-\gamma T}\Big) + \frac{1}{1+\lambda}e^{-T/\lambda}\right] = \gamma
$$

holds and $(1-p)/p$ ranges in $(0, \infty)$. Thus the sufficient condition for the transition to be discontinuous is that the second term is locally an increasing function at the origin of $\lambda$. The first order derivative of the second term at the origin is

$$\lim_{\lambda \to +0} \frac{\partial}{\partial \lambda}(1+\lambda)\frac{\partial}{\partial \lambda} \ln\left[\frac{1}{\gamma\lambda - 1}\left(e^{-T/\lambda} - e^{-\gamma T}\right) + \frac{1}{1+\lambda}e^{-T/\lambda}\right] = \gamma(1+\gamma).$$

Therefore, we can conclude that the critical probability $p_c \in (0, 1)$ exits and at which the discontinuous transition is triggered.

The rest of the section is devoted for the critical $\gamma$. Here we show that there is a value of $\gamma$ at which the optimal $\lambda$ value becomes non-zero. First, we see that there is a parameter region in which the optimal $\lambda$ is zero. $F_I(\lambda, \gamma = 0, p, T)$ is a strict monotonically decreasing function of $\lambda$, and is a continuous function of $\gamma$ for $0 < p < 1$ and $T > 0$. Thus, for a sufficiently small $\gamma > 0$, $F_I(\lambda, \gamma, p, T)$ is also the monotonically decreasing function of $\lambda$ meaning that the optimal $\lambda$ is zero.

To see there is a transition of the optimal $\lambda$ value from zero to non-zero, we derive a sufficient condition of $\lambda = 0$ being no longer optimal. Since $\gamma$ represents the killing rate of the bacterial cells, $F_I(\lambda, \gamma, p, T) \geq F_I^{\infty}(\lambda, p, T)$ and $F_I(\lambda^*, \gamma, p, T) \geq F_I^{\infty}(\lambda^*_{\infty}, p, T)$ hold regardless of the parameter values. Therefore, if $\lambda = 0$ is the optimal $\lambda$ value,

$$F_I(0, \gamma, p, T) = -pT(1+\gamma) \geq -\frac{pT}{2}\left(1+\sqrt{1+\frac{4}{pT}}\right) - 2\coth^{-1}\left(\sqrt{1+\frac{4}{pT}}\right) = F_I^{\infty}(\lambda^*_{\infty}, p, T)$$

holds. Since the inequality is the necessary condition of $\lambda = 0$ being the optimal value, the contraposition of this argument is used as the sufficient condition of the transition.

Note that $F_I(0, \gamma, p, T)$ is a monotonically decreasing function of $\gamma$ with $-\infty$ as its value at $\gamma \to \infty$ limit, whereas $F_I^{\infty}$ is the constant function. Therefore, there is a value of $\gamma$ $(= \delta)$ which satisfies $F_I(0, \delta, p, T) = F_I^{\infty}(\lambda^*_{\infty}, p, T)$. In $\gamma > \delta$ region, $F_I(0, \gamma, p, T)$ is smaller than $F_I^{\infty}(\lambda^*_{\infty}, p, T)$, i.e., $\lambda = 0$ is no longer the optimal. From the previous argument, one can see that $\delta \neq 0$ holds.

Note that $F_I(\lambda^*, \gamma, p, T) > F_I^{\infty}(\lambda^*_{\infty}, p, T)$ always holds for finite $\gamma$ values, and thus, the transition of the optimal $\lambda$ takes place at value of $\gamma$ $(= \gamma_c)$ which is strictly lower than $\delta$.

Next, we derive an upper bound of $\gamma_c$. Note that $\lambda^* = 0$ is equivalent to that the equation $\partial F_I/\partial(1/\lambda) = 0$ has no solution of $v \equiv 1/\lambda$ in $\mathrm{R}^+$. By taking the partial derivative, we get

$$\begin{aligned} \frac{\partial F_I}{\partial v} &= \frac{1}{v} - \frac{1}{1+v} + \frac{p}{\gamma - v} - p\frac{T(1+\gamma)e^{-vT} + e^{-\gamma T}}{(1+\gamma)e^{-vT} - (1+v)e^{-\gamma T}} \\ &= \frac{1 - H(v, \gamma, p, T)}{v(1+v)}, \end{aligned} \tag{12}$$

where $H(v,\gamma,p,T) \geq 0$ holds (described later) regardless of parameter values being given as

$$H(v,\gamma,p,T) = \frac{pv(1+v)(1+\gamma)\Big[e^{-\gamma T} - \big(1 - T(\gamma - v)\big)e^{-vT}\Big]}{(\gamma - v)\Big((1+\gamma)e^{-vT} - (1+v)e^{-\gamma T}\Big)}$$

Here, $H(0,\gamma,p,T) = 0$ and $\lim_{v\to\infty} H(v,\gamma,p,T) = (1+\gamma)p$ holds. Therefore, if $\gamma$ is greater than $1/p - 1$, $\partial F_I/\partial v = 0$ has a solution. This solution is only the solution if $H$ is a monotonic function of $v$, but from the existence of $\delta$, $H$ always has a solution even if $\gamma$ is smaller than $1/p - 1$. Thus, $\bar{\gamma}_c = 1/p - 1$ gives the upper bound of $\delta$ and $\gamma_c$ satisfying $\bar{\gamma}_c \geq \delta > \gamma_c$.

As shown in Fig. S7, $H$ approaches to a monotonic function as $T$ becomes smaller, whereas for large $T$ values, $H$ shows the non-monotonic feature and $\partial F/\partial v = 0$ has a solution even if $\gamma$ is smaller than $1/p - 1$.

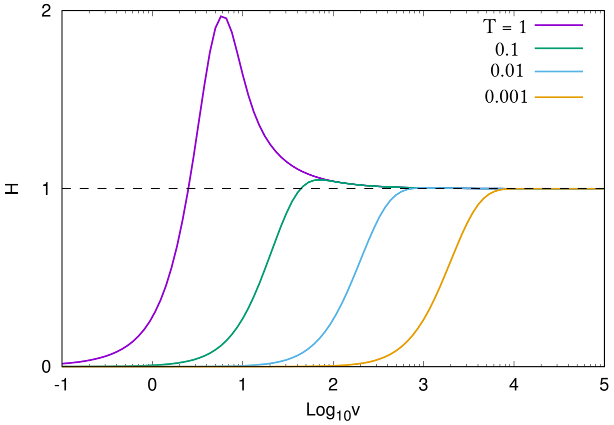

Fig.S7: **The shape of the function** $H$ $H$ is plotted as a function of $v$. $p = 0.2$ and $\gamma = 1/p - 1$.

Lastly, we show that $\lambda = 0$ has no singularity while the transition is taking place to see the transition is discontinuous. While $\partial F_I/\partial\lambda$ is not continuous at $\lambda = 0$, its right limit is given as $p(1+\gamma) - 1$. Since $0 < \gamma_c < \bar{\gamma}_c$ holds and $p(1+\gamma) - 1$ is negative in the region of $\gamma < \bar{\gamma}_c$, $\lambda = 0$ is locally optimal at $\gamma = \gamma_c$ which means that $\lambda^* > 0$ is not continuously branching out from $\lambda^* = 0$.

$H(v,\gamma,p,T) \geq 0$ is shown as follows; first, the denominator can be rewritten as

$$(\gamma - v)(1+v)e^{-\gamma T}\left(\frac{1+\gamma}{1+v}e^{(\gamma - v)T} - 1\right).$$

For the case of $\gamma > v$, $(1+\gamma)/(1+v) > 1$ and $e^{(\gamma - v)T} > 1$ hold, and thus, the sign of the large parenthesis is positive and $(\gamma - v) > 0$ which means the denominator has the positive sign. For the opposite case, $(1+\gamma)/(1+v)e^{(\gamma-v)T}$ is less than one and $(\gamma - v) < 0$. Therefore, the denominator is again positive.

The square bracket of the numerator is rewritten as

$$e^{-\gamma T}(1-(1-a)e^{a}),$$

where $a = (\gamma - v)T$. Here we introduce the function $h(a)$ which is given as $h(a) = 1-(1-a)e^{a}$. By noting that $h(0) = 1$ and $dh/da = ae^{a}$ hold, one can see that $h(a)$ monotonically increases (decreases) with $a$ in the region of $a > 0$ ($a < 0$), and $h(0)$ is the minimum. Therefore, $h(a)$ is always larger than one, and accordingly, positive.

### 8 A condition for a discontinuous transition for arbitrary probability distribution functions

In this section, we provide a proof for a sufficient condition for the sequential models with distributed antibiotics application time to exhibit the discontinuous transition of the optimal lag time. The sufficient condition is that the lower bound of the support of $q(T)$, the distribution function of the antibiotics application time, is non-zero. Note that, however, this is not a necessary condition.

For arbitrary functions $q(T)$ satisfying the condition, there exists the critical probability of the antibiotics application, $p_c$ in $(0,1)$ such that the optimal lag time is 0 for $p < p_c$ while it discontinuously transits at to a non-zero value at $p = p_c$. We denote $\mathrm{supp}(q)$ by $(T_{\mathrm{lb}}, T_{\mathrm{ub}})$. While we assume that the support consists of the single connected interval, the following argument is easily extended for cases that the support has more than one connected components.

This claim is shown as following: first, the fitness function is

$$\mathcal{F}_I = (1-p)\ln f(0) + p\langle \ln f(T)\rangle,$$

where $f(T)$ is the single-round fitness of $M$ states model given as

$$\begin{aligned} f(T) &= e^{-(1+\gamma)T}(1-\gamma\lambda)^{-(1+M)} \\ &+ e^{-(1+1/\lambda)T}\sum_{n=1}^{M+1}\frac{(T/\lambda)^{M+1-n}}{(M+1-n)!}\Big((1+\lambda)^{-n}-(1-\gamma\lambda)^{-n}\Big), \end{aligned}$$

and $\langle\cdot\rangle$ means the average over the distribution $q(T)$ (here we dropped the subscript $M$ on $\lambda$ for the readability). Accordingly, the first derivative of the fitness function is given by

$$c\frac{\partial \mathcal{F}_I}{\partial \lambda} = \frac{\partial\langle \ln f(T)\rangle/\partial\lambda}{\partial \ln f(0)/\partial\lambda} - \Big(-\frac{1-p}{p}\Big) \equiv \mathcal{G}_I(\lambda,\gamma) - \mathcal{H}_I(p),$$

where $1/c = p\cdot\partial\ln f(0)/\partial\lambda$. The non-zero optimal $\lambda$ is determined by $\mathcal{G}_I(\lambda,\gamma) = \mathcal{H}_I(p)$. Since $\mathcal{H}_I(p)$ diverges to $-\infty$ as $p \to 0$ and $\mathcal{G}_I$ is bounded, the transition triggered by $p$ happens when $\mathcal{H}(p)$ approaches to $\mathcal{G}(\lambda,\gamma)$ from below. Here, the sufficient condition for the discontinuous transition of the optimal lag time is $\mathcal{G}_I(0,\gamma) < 0$ and $\partial\mathcal{G}_I(\lambda,\gamma)/\partial\lambda|_{\lambda=0} < 0$. If $T_{\rm lb}$ is greater than zero, the calculation gives the same value for arbitrary distribution functions $q(T)$ as shown below

$$\begin{aligned}
\lim_{\lambda\to+0}\mathcal{G}_I &= \lim_{\lambda\to+0}\frac{\partial\left(\int_{T_{\rm lb}}^{T_{\rm ub}} q(T)\ln f(T)dT\right)/\partial\lambda}{\partial\ln f(0)/\partial\lambda}\\
&= \int_{T_{\rm lb}}^{T_{\rm ub}} q(T)\lim_{\lambda\to+0}\frac{\partial\ln f(T)/\partial\lambda}{\partial\ln f(0)/\partial\lambda}dT\\
&= -\gamma
\end{aligned}$$

$$\begin{aligned}
\lim_{\lambda\to+0}\frac{\partial}{\partial\lambda}\mathcal{G}_I &= \lim_{\lambda\to+0}\frac{\partial}{\partial\lambda}\frac{\partial\left(\int_{T_{\rm lb}}^{T_{\rm ub}} q(T)\ln f(T)dT\right)/\partial\lambda}{\partial\ln f(0)/\partial\lambda}\\
&= \int_{T_{\rm lb}}^{T_{\rm ub}} q(T)\lim_{\lambda\to+0}\frac{\partial}{\partial\lambda}\frac{\partial\ln f(T)/\partial\lambda}{\partial\ln f(0)/\partial\lambda}dT\\
&= -\gamma(1+\gamma)
\end{aligned}$$

Thus, the sufficient condition is fulfilled for any $q(T)$. □

Note that $T\to+0$ and $\lambda\to+0$ limit are non-communicative because

$$\lim_{\lambda\to+0}\lim_{T\to+0}\frac{\partial^n f}{\partial\lambda^n} = (-1)^n\frac{(M+n)!}{M!}$$

and

$$\lim_{T\to+0}\lim_{\lambda\to+0}\frac{\partial^n f}{\partial\lambda^n} = \gamma^n\frac{(M+n)!}{M!}$$

hold. Therefore, $T_{\rm lb} > 0$ is needed for the calculations from the fist line to the second line in above equations.

Next, we prove that the condition is not necessary by analytically showing one simple example exhibiting the discontinuous transition even though $T_{\rm lb}$ is zero. Here, we chose the number of step $M = 1$ (single-step model) and the uniform distribution in $[0,a]$ $(a > 0)$ as the distribution of the antibiotics application time $q(T)$. Then, the integral above leads to

$$\begin{aligned}
\int_0^\infty q(T)\ln f(T)dT &= \frac{1}{a}\int_0^a \ln f(T)dT\\
&= -\frac{1}{2}a\gamma - \ln(1-\gamma\lambda)\\
&+ \frac{\lambda}{a(1-\gamma\lambda)}\left[\mathrm{Li}_2\Big(\frac{1+\gamma}{1+\lambda}\lambda e^{a(\gamma-1/\lambda)}\Big) - \mathrm{Li}_2\Big(\frac{1+\gamma}{1+\lambda}\lambda\Big)\right],
\end{aligned}$$

where $\mathrm{Li}_*$ is Polylogarithm.

With this choice, $\lambda \to 0$ limits of the functions $\mathcal{G}_I$ and $\partial \mathcal{G}_I/\partial\lambda$ are given as

$$\begin{aligned}
\lim_{\lambda\to+0} \mathcal{G}_I &= -\gamma \\
\lim_{\lambda\to+0} \frac{\partial}{\partial\lambda}\mathcal{G}_I &= -(1+\gamma)(\gamma - 2/a)
\end{aligned}$$

Since $\lim_{\lambda\to+0}\mathcal{G}_I < 0$, the optimal lag time always transits from zero to finite. In addition, in this case, there is a critical value of $a$ (the domain size of $q(T)$), $a_c = 2/\gamma$, where the nature of the transition changes from continuous to discontinuous.

For $a \le a_c$, the transition is continuous, while in the other region, the optimal lag time transits discontinuously. For the confirmation of the analytic result, $a_c = 2/\gamma$, we numerically inferred the critical $a$ value. The procedure if following; under a given $\gamma$ value, we change the value of $a$. Since the optimal lag time eventually transits by increasing $p$ for any combination of $\gamma$ and $a$, we computed the optimal lag time slightly above the critical $p$ value for $(\gamma, a)$.

The optimal lag time slightly above the critical point shows discontinuous behavior as a function of $a$ (see inset of Fig. S8) indicating that the nature of the transition alters. Thus, we adopt this $a$ value as the numerically inferred critical $a$ value. As shown in Fig. S8, the numerically inferred critical $a$ and analytically-obtained critical $a$, $a_c = 2/\gamma$, collapse quite well.

Due to the difficulty of carrying out the integral, we analytically showed that the discontinuous transition takes place only for $q(T) = 1/a$, supp $q(T) = [0, a]$. However, the discontinuous transition can happen also with other distribution function, for instance, Gaussian. As shown in Fig.2 in the main text, increasing the average or the standard deviation of Gaussian makes the transition more and more steeper, and eventually, it seems that the discontinuous transitions are triggered.

For the discrete distribution $\{q_i\}_{i=0}^{N-1}$ where $q_i$ represents the probability of the antibiotics application time to be $T_i$, we can use $\sum_{i=0}^{N-1} q_i\delta(T - T_i)$ as $q(T)$. Since the lower bound of the support of $q(T)$ is now always non-zero, the optimal lag time shows the discontinuous transition as $p$ increases.

In contrast to the argument, this condition is shown also as the necessary condition for the delta function-type model described in Section.10 where the role of a non-zero lower bound of $q(T)$ is explored more in detail.

### 9 Optimal lag time distributions

#### 9.1 The general form of the optimal lag time distribution

In this section, we show that the optimal lag time distribution (without specifying details of the waking-up dynamics) has the delta function at the origin, including the case that its coefficient is zero, and a gap region next to the origin

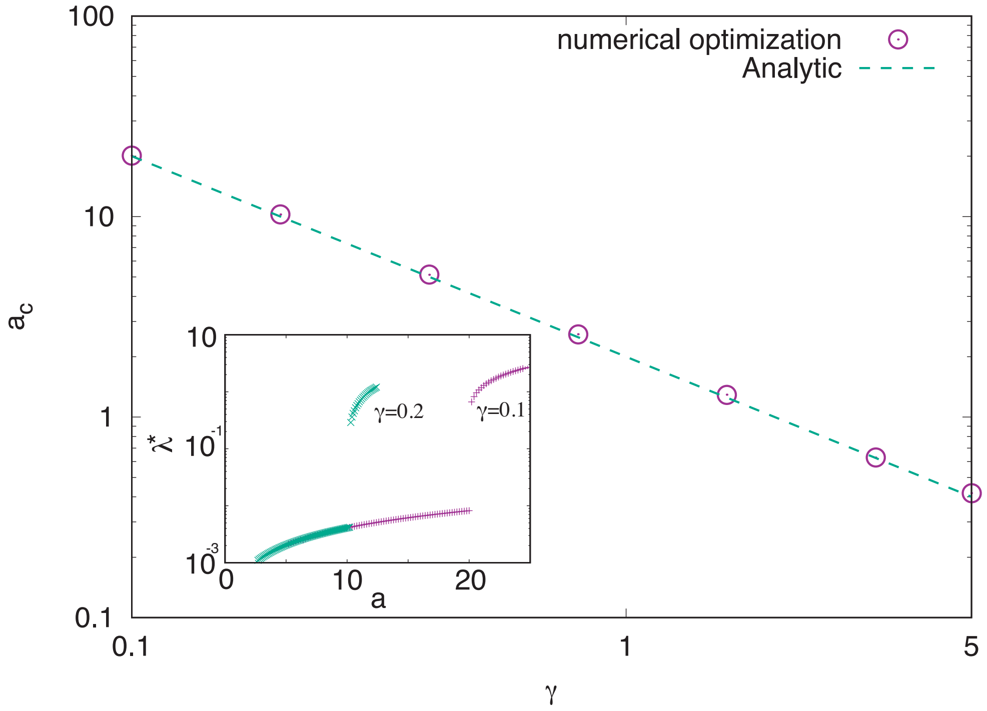

Fig.S8: **The critical a** Numerically inferred critical $a$ value for given $\gamma$ values (purple circles) are compared with analytically obtained critical $a$ value ($a_c = 2/\gamma$, green dashed line). The detailed procedure is described in the text. (inset) The optimal lag time slightly above the critical $p$ value as a function of $a$. The jump indicates that the transition of the optimal lag time at critical $p$ becomes discontinuous from continuous. For the computation, the range of $p$, $[0, 1]$, is divided into $10^4$ bins, while $10^5$ bins for $\gamma = 5.0$.

in which the probability is zero regardless of the values of $p, \gamma$, and the shape of $q(T)$.

First of all, we point out that all the piecewise continuous probability distribution functions can be decomposed as

$$r_\alpha(l) = \alpha\delta(l) + (1-\alpha)r_0(l)$$

with $\alpha \in [0,1]$, where $\delta(l)$ is the delta function peaked at the origin and $r_0(l)$ is a piecewise continuous probability distribution function satisfying $r_0(0) < \infty$. Note that $r_0(l)$ is also allowed to have at most a finite number of the delta functions.

Here we assume that $r_\alpha(l)$ is the optimal lag time distribution with $r_0(l)$ which is chosen from the set of the probability distribution functions with zero lower bound of its support[1]. We suppose $\alpha < 1$ in the following argument without losing generality because if $\alpha = 1$ is the optimal value, we can regard the support of $r_0$ as $\emptyset$ and $\mathrm{minsupp}(r_0) = \infty$.

We add an extra argument $\tau$ indicating the lower bound of the support of $r_0$ to call $r_0(l,\tau)$ as the truncated distribution function while $r_0(l,0)$ is the original $r_0(l)$. One of the natural ways to define the truncated distribution function from $r_0(l)$ is

$$r_0(l,\tau) = \begin{cases} 0 & (l < \tau) \\ r_0(l,0)/\big(\int_\tau^\infty r_0(x,0)dx\big), & (l \geq \tau) \end{cases}$$

so that $r_0(l,\tau)$ is normalized. The support can consist of more than one disconnected intervals.

To see if having a non-zero lower bound increases the fitness value, we evaluate the derivative of the average fitness $F$

$$\begin{aligned} F[r,q](\gamma,p) &= (1-p)\ln\left[\int_0^\infty e^{-l} r(l)dl\right] \\ &+ p\int_0^\infty q(T)\ln\left[e^{-(1+\gamma)T}\int_0^T e^{\gamma l} r(l)dl + \int_T^\infty e^{-l} r(l)dl\right]dT. \end{aligned}$$

with $r = r_\alpha(l,\tau)$ by $\tau$ at $\tau = 0$. Because the single-round fitness $f(T)$ is given as

$$f(T) = \begin{cases} \alpha e^{-(1+\gamma)T} + (1-\alpha)\int_\tau^\infty e^{-x}\frac{r_0(x,0)}{\int_\tau^\infty r_0(y,0)dy}dx & (T < \tau) \\ e^{-(1+\gamma)T}\Big(\alpha + (1-\alpha)\int_\tau^T e^{\gamma x}\frac{r_0(x,0)}{\int_\tau^\infty r_0(y,0)dy}dx\Big) + (1-\alpha)\int_T^\infty e^{-x}\frac{r_0(x,0)}{\int_\tau^\infty r_0(y,0)dy}dx & (T \geq \tau) \end{cases} \tag{13}$$

[1]We can construct such $r_\alpha(l)$ as following; first, find $r_0(l)$ from the set of the probability distribution function with zero lower bound of its support and without the delta function at the origin. Since $F[\alpha\delta(l) + (1-\alpha)r_0(l)] \geq F[r_0(l)]$ holds for any $r_0(l)$ including the case that $\alpha = 0$ (the case with $\alpha > 0$ will be shown later), we can choose optimal $\alpha$ for this $r_0(l)$.

the derivative is calculated as

$$\begin{aligned}
\frac{\partial F}{\partial \tau}(\tau) &= \int_0^\infty \hat{q}(T)\frac{\partial \ln[f(T)]}{\partial \tau}dT \\
&= \int_0^\tau \frac{\hat{q}(T)}{f(T)}\frac{\partial}{\partial \tau}\left[\alpha e^{-(1+\gamma)T} + (1-\alpha)\int_\tau^\infty e^{-x}\frac{r_0(x,0)}{\int_\tau^\infty r_0(y,0)dy}dx\right]dT \\
&+ \int_\tau^\infty \frac{\hat{q}(T)}{f(T)}\frac{\partial}{\partial \tau}\left[e^{-(1+\gamma)T}\Big(\alpha + (1-\alpha)\int_\tau^T e^{\gamma x}\frac{r_0(x,0)}{\int_\tau^\infty r_0(y,0)dy}dx\Big)\right. \\
&+ \left.(1-\alpha)\int_T^\infty e^{-x}\frac{r_0(x,0)}{\int_\tau^\infty r_0(y,0)dy}dx\right]dT \\
&= (1-\alpha)\int_0^\tau \frac{\hat{q}(T)}{f(T)}\left[-e^{\tau}\frac{r_0(\tau,0)}{\int_\tau^\infty r_0(y,0)dy} + \int_\tau^\infty e^{-x}\frac{\partial}{\partial \tau}\frac{r_0(x,0)}{\int_\tau^\infty r_0(y,0)dy}dx\right]dT \\
&+ (1-\alpha)\int_\tau^\infty \frac{\hat{q}(T)}{f(T)}\left[-e^{-T-\gamma(T-\tau)}\frac{r_0(\tau,0)}{\int_\tau^\infty r_0(y,0)dy} + \int_\tau^T e^{\gamma x}\frac{\partial}{\partial \tau}\frac{r_0(x,0)}{\int_\tau^\infty r_0(y,0)dy}dx\right. \\
&+ \left.\int_T^\infty e^{-x}\frac{\partial}{\partial \tau}\frac{r_0(x,0)}{\int_\tau^\infty r_0(y,0)dy}dx\right]dT
\end{aligned}$$

with $\hat{q}(T)$ as $(1-p)\delta(T) + pq(T)$. From the definition of $r_0(x,\tau)$,

$$\frac{\partial}{\partial \tau}r_0(x,\tau) = \frac{r_0(\tau,0)r_0(x,0)}{\left(\int_\tau^\infty r_0(y,0)dy\right)^2} \to r_0(0,0)r_0(x,0), \text{ (as } \tau \to 0)$$

holds. By substituting $\tau = 0$ to the equation above, we get

$$\begin{aligned}
c\frac{\partial F}{\partial \tau}\bigg|_{\tau=0} &= \int_0^\infty \frac{\hat{q}(T)}{f(T)}\left(e^{-(1+\gamma)T}\int_0^T e^{\gamma x}r_0(x,0)dx + \int_T^\infty e^{-x}r_0(x,0)dx\right)dT \\
&- \int_0^\infty \frac{\hat{q}(T)}{f(T)}e^{-(1+\gamma)T}dT \qquad (14)
\end{aligned}$$

where $c = ((1-\alpha)r_0(0,0))^{-1}$ $(0 < c < \infty)$. To judge whether a non-zero lower bound of the support is advantageous, we want to see the size relationship between the first and the second term in the right hand side of the equation.

Now we calculate the functional variation of $F$ by $r_\alpha$. Consider the situation that $r_\alpha(x)$ is perturbed by an arbitrary probability distribution function with zero lower bound of the support, $\eta(x)$, as $r_\alpha(x) \to r'_\alpha(x) \propto r_\alpha(x) + \epsilon\eta(x)$. Since $r'_\alpha(x)$ still needs to be a probability distribution function, it must be normalized and $\epsilon$ has to be positive. Since we chose $\eta(x)$ from the probability distribution functions, $r'_\alpha(x)$ is given as $(r_\alpha(x) + \epsilon\eta(x))/(1+\epsilon)$. By taking the derivative of

$F$ with $r'_\alpha(x)$ by $\epsilon$ at $\epsilon = 0$, we get

$$\begin{aligned}\left.\frac{\partial F}{\partial \epsilon}\right|_{\epsilon=0} &= \int_0^\infty \frac{\hat{q}(T)}{f(T)}\left(e^{-(1+\gamma)T}\int_0^T e^{\gamma x}(\eta(x)-r_\alpha(x))dx\right. \\ &+ \left.\int_T^\infty e^{-x}(\eta(x)-r_\alpha(x))dx\right)dT \qquad (15)\end{aligned}$$

Note that $f(T)$ is calculated with $r_\alpha(x)$, but not $r'_\alpha(x)$ because $\epsilon = 0$ is substituted. Because of the optimality, $\partial F/\partial\epsilon$ at $\epsilon = 0$ needs to be smaller than or equals to zero. This is not necessarily the extreme point as usual functional variations lead because the perturbation with $\epsilon < 0$ is not allowed in this problem. However, if it is zero, the functional $I$ defined by

$$I[a](T) = e^{-(1+\gamma)T}\int_0^T e^{\gamma x}a(x)dx + \int_T^\infty e^{-x}a(x)dx$$

with $a$ as a function with zero lower bound of its support needs to satisfy the equality

$$I[\eta_0](T) = I[r](T) = I[\eta_1](T)$$

for $^\forall T \geq 0$ where $\eta_0$ and $\eta_1$ are different probability distribution function because $q(T)$ is an arbitrary function. By choosing, for instance, an exponential distribution with the parameter $\lambda_i$ ($\lambda_0 \neq \lambda_1$) as $\eta_i(x)$, we can easily show that $I[\eta_0] \neq I[\eta_1]$. Thus, $\partial F/\partial\epsilon$ needs to be smaller than zero.

By choosing the delta function $\delta(x)$ as $\eta(x)$ the optimal condition (right hand side of Eq.(15) is smaller than zero) leads to

$$\int_0^\infty \frac{\hat{q}(T)}{f(T)}\left(e^{-(1+\gamma)T}\int_0^T e^{\gamma x}r_0(x,0)dx + \int_T^\infty e^{-x}r_0(x,0)dx\right)dT > \int_0^\infty \frac{\hat{q}(T)}{f(T)}e^{-(1+\gamma)T}dT.$$

This is what we desired. From Eq.(14) and the inequality above, $\partial F/\partial\tau\big|_{\tau=0}$ is shown to be positive, and thus, the non-zero lower bound increases the fitness. (Note that it is possible that $r_0(l,\tau)$ is suboptimal among all the function with its lower bound of the support as $\tau$, while still it performs better than any distribution function with zero lower bound of the support.).

Therefore, if $r_\alpha$ is the optimal distribution, there is always a region next to the origin in which $r_\alpha$ is zero. □

So far, we have assumed that $0 < \alpha < 1$ can be optimal, but the existence of the optimal $\alpha$ in $(0, 1)$ is not yet shown. The remaining part of this subsection is devoted for showing it.

We evaluate the first-order derivative of $F[r_\alpha]$ by $\alpha$ at $\alpha = 0$, given as

$$\frac{\partial F}{\partial \alpha} = \int_0^\infty \hat{q}(T)\frac{\partial}{\partial\alpha}\ln[f(T,\alpha)]dT$$

where we added $\alpha$ as an argument of $f$ to emphasize that $f$ is a function of $\alpha$. The existence of $\alpha$ satisfying of $\partial F/\partial\alpha = 0$ in $[0, 1]$ is equivalent to that the optimal $\alpha$ is in $[0, 1]$, and otherwise, the optimal $\alpha$ is either zero or unity. Therefore, if we show the uniqueness of the solution in $[0, 1]$ for given $(\gamma, p, q(T))$, the continuity of the optimal $\alpha$ on $(\gamma, p, q(T))$ will be shown because the derivative is a continuous function of $\gamma, p$, and $q(T)$ (for $q(T)$, we need to calculate the functional variation of $F$). By noting that $f$ is linear in $\alpha$, $\partial_\alpha f$ is a constant function in $\alpha$. Therefore, the second order derivative is given as

$$\frac{\partial^2 F}{\partial\alpha^2} = -\int_0^\infty \hat{q}(T)\frac{(\partial_\alpha f)^2}{f^2}dT < 0$$

meaning that $\partial_\alpha F$ is a monotonically decreasing function of $\alpha$, and thus, there is at most one solution in $[0, 1]$. Now the optimal $\alpha$, $\alpha^* = \alpha^*(\gamma, p)[q]$ is a continuous function.

$\alpha^*(\gamma, 0)[q] = 1$ holds because of the monotonicity of $e^{-x}$. Also, there is the critical values of $p$ and $\gamma$ at which the optimal $\alpha$ transits to zero if the delta function is chosen as $q(T)$ (see Section.10). Thus, there is a way to change the optimal $\alpha$ continuously from unity to an arbitrary value by continuously modulating $(p, \gamma, q(T))$.

#### 9.2 Computing the optimal lag time distribution

##### 9.2.1 Derivation of the optimal condition

In this section, we present the method to obtain the optimal lag time distribution for a given distribution of the antibiotics application time, $q(T)$. If the distribution is discrete, given as $\mathrm{Prob}(T_i) = q_i$, $(T_i > 0)$ for $i = 1, \cdots, M-1$ and $\sum_{i=1}^{M-1} q_i = 1$, we introduce the expression $q(T) = \sum_{i=1}^{M-1} q_i\delta(T - T_i)$ for the unified description.

Here, we compute the optimal lag time distribution for $q(T)$ with the finite support $[0, T_{\max})$ while the case for the infinite support can be considered by taking $T_{\max} \to \infty$ limit [2]. Then, the upper bound of the optimal lag time distribution becomes $T_{\max}$ because it just decreases the fitness to invest any fraction of population to have lag times longer than $T_{\max}$. (For the effect of $T_{\max}$ to the optimal lag time, see Fig.S9).

Now the fitness function is given as

$$F = \int_0^{T_{\max}} \hat{q}(T)\ln\Big[\int_0^{T_{\max}} h(T, l)r(l)dl\Big]dT$$

where $\hat{q}(T) = (1-p)\delta(T) + pq(T)$ and $h(T, l)$ is

$$h(T, l) = \begin{cases}\exp[-T - \gamma(T - l)] & (l < T)\\ \exp[-l] & (\text{otherwise}).\end{cases} \tag{16}$$

[2] Note that the choice of $T_{\max}$ corresponds to selecting a way of the antibiotics application. Even if two distribution functions $q_1(T)$ and $q_2(T)$ with the support $[0, T_{\max}^{(1)})$ and $[0, T_{\max}^{(2)})$, respectively, converge to the same distribution $q(T)$ in $T_{\max}^{(1)}, T_{\max}^{(2)} \to \infty$ limit, the optimal lag time distributions for $q_1(T)$ and $q_2(T)$ are different in general.

Next, we approximate the optimal lag time distribution by discretizing it with the bins of size $\Delta$, $\{r_n\}_{n=0}^{N-1}$ where $N = T_{\max}/\Delta$. Now we obtain

$$F_d(T) = \int_0^{T_{\max}} \hat{q}(T) \ln\Big[\sum_{n=0}^{N-1} h(T, n\Delta) r_n\Big] dT.$$

Owing to the discretization, now we can carry out the maximization of the function $F_d$ by taking differentials of $F_d$ respect to $r_n'$s. Since $r_n'$s are the discrete probability distribution function, $r_n \geq 0$ and $\sum_{n=0}^{N-1} r_n = 1$ needs to be satisfied.

This can be solved by introducing KKT multipliers $\mu_n$ for the condition $r_n \geq 0$ and $\bar{\mu}$ for the condition $\sum_{n=0}^{N-1} r_n = 1$ as

$$L(\vec{r}, \vec{\mu}) = \int_0^{T_{\max}} \hat{q}(T) \ln\left[\sum_{n=0}^{N-1} h(T, n\Delta) r_n\right] dT + \sum_{n=0}^{N-1} \mu_n r_n - \bar{\mu} \sum_{n=0}^{N-1} r_n,$$

and imposing

$$\frac{\partial L}{\partial r_n} = 0, \;\; \mu_n r_n = 0, \;\; \mu_n \geq 0, \;\; r_n \geq 0, \;\; (0 \leq n < N), \sum_{n=0}^{N-1} r_n = 1.$$

This results in

$$\bar{\mu} - \mu_n = \int_0^{T_{\max}} \hat{q}(T) \frac{h(T, n\Delta)}{\langle h(T) \rangle} dT \tag{17}$$

$$r_n = 0 \quad \text{or} \quad \mu_n = 0. \tag{18}$$

with $r_n \geq 0$ and $\mu_n \geq 0$ for $n = 0, 1, \cdots, N-1$. $\langle h(T) \rangle$ represents the mean, i.e., $\langle h(T) \rangle = \sum_{n=0}^{N-1} h(T, n\Delta) r_n$. The Lagrange multiplier $\bar{\mu}$ is determined by multiplying eq. (17) with $r_n$ and taking the sum over $n$. Knowing $r_n \mu_n = 0$, this leads to

$$\bar{\mu} \left(\sum_{n=0}^{N-1} r_n\right) = \int_0^{T_{\max}} \hat{q}(T) \frac{\sum_{n=0}^{N-1} r_n h(T, n\Delta)}{\langle h(T) \rangle} dT = \int_0^{T_{\max}} \hat{q}(T) dT = 1,$$

giving $\bar{\mu} = 1$.

Next, we carry out the integral over $T$. If the distribution of $T$ is discrete, the integral leads to

$$\int_0^{T_{\max}} \hat{q}(T) \frac{h(T, n\Delta)}{\langle h(T) \rangle} dT = \sum_{m=0}^{M-1} \hat{q}_m \frac{h(T_m, n\Delta)}{\langle h(T_m) \rangle},$$

where $\hat{q}_0 = (1-p)$ and $\hat{q}_m = p q_m$ $(m \geq 1)$.

If $q(T)$ is a continuous function, since $h(T, l)$ has different expression depending on the size relationship between $T$ and $l$, we divide the integral into parts so that $h(T, l)$ has the same expression in each integral as follows;

$$\int_0^{T_{\max}} \hat{q}(T) \frac{h(T, n\Delta)}{\langle h(T) \rangle} dT = \sum_{m=0}^{N-1} \int_{m\Delta}^{(m+1)\Delta} \hat{q}(T) \frac{h(T, n\Delta)}{\langle h(T) \rangle} dT. \tag{19}$$

Then, we assume that $\Delta$ is sufficiently small so that each integral is well-approximated by

$$\hat{q}_m \frac{h(m\Delta, n\Delta)}{\langle h(m\Delta)\rangle},$$

where $\hat{q}_m$ is $\Delta \cdot \hat{q}(m\Delta)$ for $m \neq 0$ while $\hat{q}_0$ by $(1-p)+pq(0)\Delta$. Now, by rewriting $h(m\Delta, n\Delta)$ and $\langle h(m\Delta)\rangle$ (or $h(T_m, n\Delta)$ and $\langle h(T_m)\rangle$ in the discrete case) as $h_n^m$ and $\langle h^m\rangle$, the condition for the $n$th bin is given as

$$1 - \mu_n = \sum_{m=0}^{S-1} \hat{q}_m \frac{h_n^m}{\langle h^m\rangle},\ (r_n = 0 \text{ or } \mu_n = 0) \tag{20}$$

with $r_n \geq 0$ and $\mu_n \geq 0$, where $S = N$ for a continuous $q(T)$ while $M$ for discrete $q(T)$.

A remarkable feature of the equations is that the distribution $\{r_i\}_{i=0}^{N-1}$ appears only in the form of the averages and the equations are independent linear equations in $1/\langle h^m\rangle$. Because of the linearity of the equations, as many equalities as the number of terms in the sum are satisfied. If $q(T)$ is the continuous function, the number of non-zero $\hat{q}_m'$s equals to $N$, and thus, all $N$ equations are solvable while it is not guaranteed that the solution satisfies the constraint $r_i \geq 0$. In contrast, if $q(T)$ is a discrete function, the number of averages appearing in the sum can be either less or more than $N$. However, if we take $\Delta$ sufficiently small, $M < N$ generally holds. Hereafter, we suppose $q$ is a continuous distribution function, and accordingly, the number of non-zero $\hat{q}_m'$s is $N$.

It is proven in the previous section that the optimal lag time distribution generally has a gap region next to the origin. Thus, some $r_n'$s need to be set to zero meaning that we cannot use the equation (Eq.(20)) for some $n'$s to determine the optimal lag time distribution. However, it is worth mentioning that if we virtually remove the constraints $r_n \geq 0$ (then $\mu_n = 0$ is the solution for all $\mu_n'$s), we can solve $r_n'$s via solving the average $\langle h^m\rangle$.

By denoting $1/\langle h^m\rangle$ by $x_m$ and supposing $x_m's$ as variables, Eq.(20) is regarded as a system of linear equations, and thus, the solution is given as $\mathbf{x} = A^{-1}\mathbf{1}$ where

$$A = \begin{pmatrix} \hat{q}_0 h_0^0 & \hat{q}_1 h_0^1 & \dots & \hat{q}_{N-1} h_0^{N-1} \\ \hat{q}_0 h_1^0 & \hat{q}_1 h_1^1 & \dots & \hat{q}_{N-1} h_1^{N-1} \\ \vdots & \vdots & \ddots & \vdots \\ \hat{q}_0 h_{N-1}^0 & \hat{q}_1 h_{N-1}^1 & \dots & \hat{q}_{N-1} h_{N-1}^{N-1} \end{pmatrix},\quad \mathbf{1} = \begin{pmatrix} 1 \\ 1 \\ \vdots \\ 1 \end{pmatrix}.$$

From the definition of the average, the lag time distribution $\{r_i\}_{i=0}^{N-1}$ are given as $\mathbf{r} = B^{-1}\mathbf{x}_{\text{inv}}$, where

$$B = \begin{pmatrix} h_0^0 & h_1^0 & \dots & h_{N-1}^0 \\ h_0^1 & h_1^1 & \dots & h_{N-1}^1 \\ \vdots & \vdots & \ddots & \vdots \\ h_0^{N-1} & h_1^{N-1} & \dots & h_{N-1}^{N-1} \end{pmatrix},\quad \mathbf{x}_{\text{inv}} = \begin{pmatrix} 1/x_0 \\ 1/x_1 \\ \vdots \\ 1/x_{N-1} \end{pmatrix}.$$

##### 9.2.2 Solving the equations

As discussed, some of the conditions are not fulfilled without setting $r_i$ to zero. Therefore, the optimal lag time distribution is never obtained by solving the linear equation above. Instead, we need to set some $r_i$'s to zero and solve the equations. Since the $i$th equation,

$$1 - \mu_i = \sum_{j=0}^{N-1} \hat{q}_j \frac{h_i^j}{\langle h^j \rangle}$$

is no longer the condition to determine the optimal distribution then, the system equations to determine the average is now underdetermined. Thus, we first solve the average and the probability as the function of undetermined averages, and consequently, solve the self-consistent equations for the undetermied averages.

We re-order indices so that $r_i'$s with $i \in \{L \cdots N-1\} \equiv K$ are set to zero, and leave $x_i$ with $i \in K$ as free parameters to solve $x_i$ with $i \in \{0, 1, \cdots, L-1\}$ (There is no criterion to find the index set before solving the equations). We refer the $L$ by $L$ submatrix of $A$ by $A_L$. Now, $N$ linear equation with $N$ variable becomes $L$ equations with $L$ variable given as

$$\begin{pmatrix} a_{00} & a_{01} & \cdots & a_{0(L-1)} \\ a_{10} & a_{11} & \cdots & a_{1(L-1)} \\ \vdots & \vdots & \ddots & \vdots \\ a_{(L-1)0} & a_{(L-1)1} & \cdots & a_{(L-1)(L-1)} \end{pmatrix} \begin{pmatrix} x_0 \\ x_1 \\ \vdots \\ x_{L-1} \end{pmatrix} = \begin{pmatrix} 1 - \sum_{i=L}^{N-1} a_{0i} x_i \\ 1 - \sum_{i=L}^{N-1} a_{1i} x_i \\ \vdots \\ 1 - \sum_{i=L}^{N-1} a_{(L-1)i} x_i \end{pmatrix}$$

where $a_{ij}$ is the $(i,j)$th element of the original $N$ by $N$ matrix $A$ after re-ordering indices. The solutions are given as

$$x_n = \Xi_n - \sum_{j=L}^{N-1} \xi_{ni} x_i, \ (0 \le n < L),$$

where $\Xi_n = \sum_{i=0}^{L-1} (A_L^{-1})_{ni}$ and $\xi_{ni} = \sum_{j=0}^{L-1} (A_L^{-1})_{nj} a_{ji}$. Similarly, we use the submatrix of $B$, $B_L$ to solve

$$\begin{pmatrix} b_{00} & b_{01} & \cdots & b_{0(L-1)} \\ b_{10} & b_{11} & \cdots & b_{1(L-1)} \\ \vdots & \vdots & \ddots & \vdots \\ b_{(L-1)0} & b_{(L-1)1} & \cdots & b_{(L-1)(L-1)} \end{pmatrix} \begin{pmatrix} r_0 \\ r_1 \\ \vdots \\ r_{L-1} \end{pmatrix} = \begin{pmatrix} 1/x_0 \\ 1/x_1 \\ \vdots \\ 1/x_{L-1} \end{pmatrix}.$$

Now $r_i' s (0 \le i < L)$ are solved as functions of undetermined averages, $x_L, x_{L+1}, \cdots, x_{N-1}$. As the last step, we need to consistently determine the averages $x_L, x_{L+1}, \cdots, x_{N-1}$. The equations is given as

$$\begin{pmatrix} b_{L0} & b_{L1} & \cdots & b_{L(L-1)} \\ b_{(L+1)0} & b_{(L+1)1} & \cdots & b_{(L+1)(L-1)} \\ \vdots & \vdots & \ddots & \vdots \\ b_{(N-1)0} & b_{(N-1)1} & \cdots & b_{(N-1)(L-1)} \end{pmatrix} \begin{pmatrix} r_0(x_L, x_{L+1}, \cdots, x_{N-1}) \\ r_1(x_L, x_{L+1}, \cdots, x_{N-1}) \\ \vdots \\ r_{L-1}(x_L, x_{L+1}, \cdots, x_{N-1}) \end{pmatrix} = \begin{pmatrix} 1/x_L \\ 1/x_{L+1} \\ \vdots \\ 1/x_{N-1} \end{pmatrix}.$$

Note that here the matrix in the left hand side of the equation is $(N-L)$ by $L$, but not square matrix. Therefore, we cannot invert the matrix, but instead, derive the self-consistency equations as follows;

$$x_{L+n} \sum_{i=0}^{L-1} \frac{\tilde{\eta}_{ni}}{1 - \sum_{j=L}^{N-1} \tilde{\xi}_{ij} x_j} - 1 = 0 \tag{21}$$

with $\tilde{\xi}_{ij} = \Xi_i^{-1} \xi_{ij}$ and $\tilde{\eta}_{ij} = \Xi_j^{-1} \sum_{k=0}^{L-1} b_{(i+L)k} (B_L^{-1})_{kj}$. By solving the system of $L$th order equation, we obtain the solutions of $x_i's$ with $i \in K$ and accordingly remaining $x_i's$ and the lag time distribution $\{r_i\}_{i=0}^{N-1}$. While Eq.(21) is not solvable symbolically, the solution of $\mathbf{x}$ obtained by solving the linear equations above works well as the initial guess of numerical solvers.

##### 9.2.3 Computational protocols

There is no criterion to find the indices which cannot satisfy the conditions without setting $r_i = 0$, and thus, in principle, only the way to obtain the optimal distribution $\{r_i\}_{i=0}^{N-1}$ is trying all the possible combination of the indices that $r_i$ is set to zero, check if all $r_j'$s which are not initially set to zero are positive, and compare the fitness of all those consistent solutions.

It is practically impossible. Thus, we take advantage of the proven form of the optimal lag time distribution. Recall that the optimal lag time has a form $u(l) = \alpha\delta(l) + (1-\alpha)u_0(l)$ with minsupp$(u_0) > 0$. Thus, we need to know how many connected components configure the support of $u_0(l)$. If the number of connected components is just one, we can carry out the computation to check all the possible zero/non-zero allocation for $r_i'$s. While this is nothing but a heuristic, we have performed numerical optimizations of the fitness function by $r_i$ values to guess the number of the connected components. The optimization indicated that the number of the connected components is the same number with the peaks of $q(T)$. Therefore, we computed all the possible distributions with a single connected support of $r_i$ $(i \geq 2)$[3]. The optimal distributions for a normal, exponential, and power-law distribution are computed in this way. This method guarantees that the obtained distribution has the highest fitness among all the locally optimal distributions whose support has two elements (origin and the support of $u_0(l)$).

Still, too much intensive computation is required if the number of connected components of supp$(u_0)$ is more than one. Thus, for the computation of the

[3]We checked only for $i \geq 2$ because $i = 0$ is for the delta function and at least $i = 1$ is for a gap as long as the discretization is done well to approximate the distributions

optimal lag time distribution for a double-Gaussian distribution shown in the main text, we adopted an iterative solving technique. The scheme is following:

(0) solve the linear equation to obtain $\mathbf{r} = B^{-1}\mathbf{x}_{\mathrm{inv}}$ add the indices $n$ to $K$ if $r_n$ becomes smaller than or equal to zero in the solution.

(1) solve the consistency equation

$$x_{L+m} \sum_{i=0}^{L-1} \frac{\tilde{\eta}_{mi}}{1 - \sum_{j=L}^{N-1} \tilde{\xi}_{ij} x_j} - 1 = 0$$

to obtain $r_n'$s, where $L + m \in K$ $(m = 0, 1, \cdots \#K - 1)$.

(2) if all the bin $r_n'$s are greater than or equal to zero, then end the iteration, otherwise, put the indices $n$ whose $r_n$ are negative to $K$ and return to (1).

The definition of $K$ is given in the second paragraph of this section, and at every update of $K$, the indices are re-ordered so that $K$ is written as $\{L, \cdots, N-1\}$. It is possible that the obtained distribution by this method is inferior to other distributions even if those distributions have the same or less numbers of the elements of the support. However, it is confirmed that this iterative method gives the same distribution led by the former method for normal and exponential distributions, but not power-law distributions as far as we have tried.

### 10 The delta function-type model

We can also consider the simplest model where the cells start to grow immediately after the time $\lambda$ has passed without any variation, i.e., the lag time distribution follows the Dirac's delta function. Since the model allows us to extract the mathematically clear essences of the population dynamics of waking-up, in this section we repeat the same analysis done for the sequential models. Some results obtained from the delta function-type model differ from the outcome of the sequential model even qualitatively. Still, it provides useful insights to us for understanding the results described above and in main text.

#### 10.1 The optimal lag time in a single, and double phenotypes case

Therefore, the temporal evolution of the bacterial population at time $t$ ($t > \max\{\lambda, T\}$) given as following (Table.1),

Then, the fitness function is given by

$$F_I^\delta(\lambda, \gamma, p, T) = \begin{cases} -p(T-\lambda)(1+\gamma) - \lambda & (\lambda < T) \\ -\lambda & (\lambda > T). \end{cases} \tag{22}$$

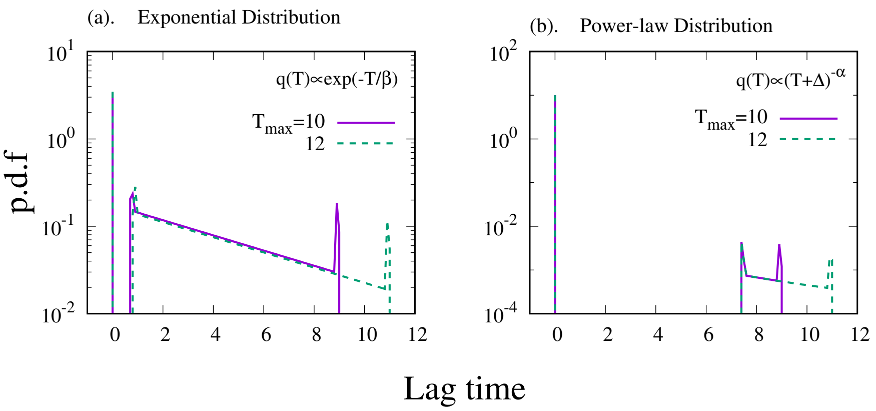

Fig.S9: **The optimal distribution with different** $T_{\max}$ The optimal distribution is computed for (a). the exponential and (b) the power-law distribution, respectively, with two different values of $T_{\max}$. Here, we compare the optimal distributions with $T_{\max} = 10$ and 12. While the exponential distribution and the power-law distribution led to the distinct optimal distributions for the different $T_{\max}$ values, the optimal distributions for the normal distribution and the sum of the two normal distributions (panel (a) and (d) in Fig.5 in the main text) were unchanged. The other parameter values are the same with Fig.5 in the main text.

| | + AB (prob. $p$) | - AB (prob. $1-p$) |
|---|---|---|
| $\lambda < T$ | $\exp[-\gamma(T-\lambda)]\exp[t-T]$ | $\exp[t-\lambda]$ |
| $\lambda > T$ | $\exp[t-\lambda]$ | $\exp[t-\lambda]$ |

Table 1: The population of the cells at time $t > \max\{\lambda, \mathrm{T}\}$ for each condition.

By taking the first-order derivative of $F_I^\delta$ respect to $\lambda$, we obtain

$$\frac{\partial F_I^\delta}{\partial \lambda}(\lambda, \gamma, p, T) = \begin{cases} p(1+\gamma)-1 & (\lambda < T) \\ -1 & (\lambda > T). \end{cases} \tag{23}$$

Therefore, the optimal lag time, $\lambda^*$ shows the discontinuous transition as

$$\lambda^* = \begin{cases} 0 & (\gamma < 1/p - 1) \\ T & (\gamma \geq 1/p - 1). \end{cases} \tag{24}$$

It is worth noting that the transition point is determined only by $\gamma$ and $p$. In contrast to the models described above and in the main text. The antibiotics application time $T$ has no role in the transition. Also, the $\gamma = 1/p - 1$ is not upper bound of the transition point but exactly determines the transition point.

The calculations for the two-strategy case are also carried out analytically. For the two species case, the optimal parameter values are given as $\lambda_a = 0$, $\lambda_b = T$, and

$$x^* = \frac{p(1-\exp[-(1+\gamma)T]) - \exp[-\gamma T](1-\exp[-T]))}{(1-\exp[-T])(1-\exp[-\gamma T])}.$$

The optimal $x$ is shown in Fig. S10 which has qualitatively the same features of the optimal fraction of the model described in the main text.

#### 10.2 The steepness of the transition and the support of $q(T)$

We have shown that a non-zero lower bound of the support of $q(T)$ is a sufficient condition for the sequential model to exhibit the discontinuous transition of the optimal lag time. To see how the lower bound of the support changes nature of the transition, here we explicitly calculate the first order derivative of the fitness by the lag time $\lambda$.

Here, a normal distribution with the average $\tau$ and the standard deviation $\sigma$ is chosen as the distribution function of $T$. Then, the fitness function is given in the form $\mathcal{F}_I^\delta = -(1-p)\lambda + p\mathcal{G}_I^\delta(\lambda, \gamma)$ and $\mathcal{G}_I^\delta$ has three different expressions depending on the size relationship among $T_{\mathrm{lb}}, T_{\mathrm{ub}}$, and $\lambda$ which are given as

$$\mathcal{G}_I^\delta = \begin{cases} \int_{T_{\mathrm{lb}}}^{T_{\mathrm{ub}}} (\gamma\lambda - (1+\gamma)T) q(T) dT & (\lambda \leq T_{\mathrm{lb}}) \\ -\int_{T_{\mathrm{lb}}}^{\lambda} \lambda q(T) dT + \int_{\lambda}^{T_{\mathrm{ub}}} (\gamma\lambda - (1+\gamma)T) q(T) dT & (T_{\mathrm{lb}} < \lambda \leq T_{\mathrm{ub}}) \\ -\int_{T_{\mathrm{lb}}}^{T_{\mathrm{ub}}} \lambda q(T) dT & (T_{\mathrm{ub}} < \lambda). \end{cases}$$

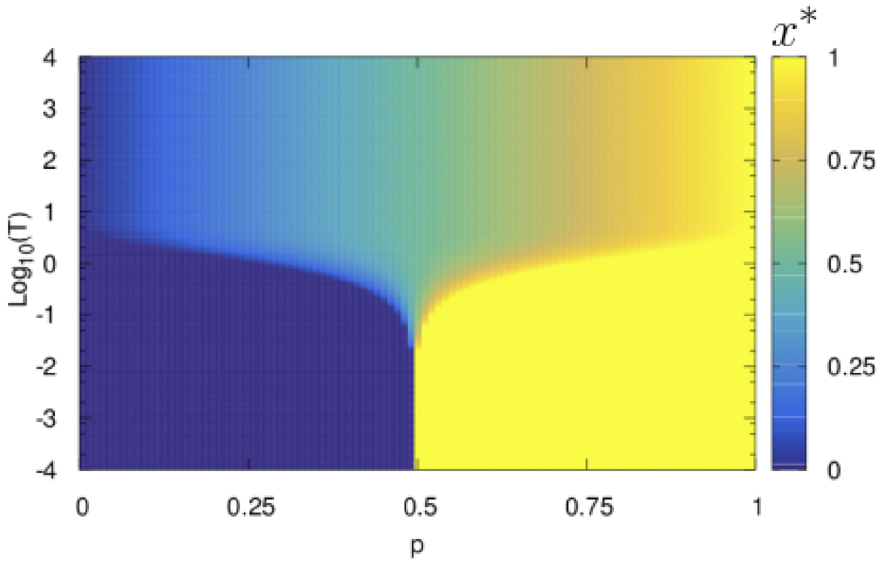

Fig.S10: **The the optimal fraction** $x^*$ The the optimal fraction $x^*$ is plotted as a function of $p$ and $T$. The steep transition at $p \approx 0.5$ in smaller $T$ region is continuous. $\gamma$ is set to unity.

In the third case, the integral simply leads to $-\lambda$ which means $\partial_\lambda \mathcal{F}_I^\delta = -1$. Thus $\lambda > T_{\mathrm{ub}}$ never be an optimal solution and there is no need to consider this region.

For the first case, by carrying out the integral, we obtain $\mathcal{G}_I^\delta = \gamma\lambda - \mathcal{H}_I^\delta(\gamma)$ where $\mathcal{H}_I^\delta$ is constant in $\lambda$[4]. Therefore, the first-order derivative of $\mathcal{F}_I^\delta$ is given as

$$\frac{\partial \mathcal{F}_I^\delta}{\partial \lambda} = -(1-p) + p\gamma.$$

The derivative shows that the optimal lag time $\lambda^*$ is zero as long as $\gamma < (1-p)/p$ holds and it transits to a some value greater than $T_{\mathrm{lb}}$.

To see what happens if the lower bound of the support is zero, here we set $T_{\mathrm{lb}} = 0$. Also, for the simplicity, we assume $T_{\mathrm{ub}} \to \infty$. This assumption has no effect to the result because the derivative of the fitness function is always minus one in the third region $\lambda > T_{\mathrm{ub}}$.

[4]It is given as

$$\mathcal{H}_I^\delta = \frac{\sqrt{2}\sigma(1+\gamma)}{\Omega}\Big((e^{-(\tau'-T_{\mathrm{lb}}')^2} - e^{-(\tau'-T_{\mathrm{ub}}')^2})/\sqrt{\pi} + \tau'(\mathrm{erf}(\tau'-T_{\mathrm{lb}}') - \mathrm{erf}(\tau'-T_{\mathrm{ub}}'))\Big),$$

where $\tau', T_{\mathrm{lb}}', T_{\mathrm{ub}}'$, and $\Omega$ is $\tau/(\sqrt{2}\sigma)$, $T_{\mathrm{lb}}/(\sqrt{2}\sigma)$, $T_{\mathrm{ub}}/(\sqrt{2}\sigma)$, and $\mathrm{erf}(\tau'-T_{\mathrm{lb}}') - \mathrm{erf}(\tau' - T_{\mathrm{ub}}')$, respectively.

Since now the support is $\mathbf{R}^+$, $\mathcal{F}_I^\delta$ has only one expression that

$$\frac{\Omega}{\sqrt{2}\sigma p}\mathcal{F} = -\lambda' \frac{1-p}{p}(1+\mathrm{erf}(\tau'))$$
$$+\lambda'(\gamma - \mathrm{erf}(\tau')) - (1+\gamma)\Big[(\lambda'-\tau')\,\mathrm{erf}(\lambda'-\tau') + \tau' + \frac{e^{-(\lambda'-\tau')^2}}{\sqrt{\pi}}\Big],$$

where the variables with $'$ represent the original variables divided by $\sqrt{2}\sigma$ and $\Omega = 1 + \mathrm{erf}(\tau')$. Then, the first-order derivative by $\lambda$ (note that it is not by $\lambda'$) is

$$\frac{\Omega}{p(1+\gamma)}\frac{\partial \mathcal{F}}{\partial \lambda} = -\,\mathrm{erf}(\lambda'-\tau') + h(\tau',\gamma,p) \tag{25}$$

where $h$ is given as

$$h(\tau',\gamma,p) = \frac{1}{1+\gamma}\Big(\gamma - \mathrm{erf}(\tau') - \frac{1-p}{p}(1+\mathrm{erf}(\tau'))\Big).$$

Let us consider the transition triggered by an increase of $p$. For the simplicity, we assume $\tau' \gg 1$ and $\gamma = 1$ leading to $h \approx -(1-p)/p$. Since $h$ diverges to $-\infty$ as $p \to 0$, the optimal lag time is zero at $p \approx 0$. As $p$ increases, $h$ approaches to $\mathrm{erf}(\lambda'-\tau')$ from below. By noting that the error function erf is the monotonically increasing function of the argument, there is only one intersection of $\mathrm{erf}(\lambda'-\tau')$ and $h$, and it is first made at $\lambda' = 0$. Therefore, the transition is continuous.

Note that $\lambda'$ and $\tau'$ are now scaled by the standard deviation of Normal distribution, $\sigma$, and thus, the flatness of $\mathrm{erf}(\lambda'-\tau')$ in $\lambda' \ll \tau'$ (and also $\lambda' \gg \tau'$) and the steepness of its sigmoidal shape depends on $\sigma$. If the distribution is greatly broad ($\sigma \gg \tau$), the error function is approximately a linear function even around $\lambda = 0$. With such broad distribution, the transition looks continuous.

However, as $\sigma$ becomes smaller and smaller, the error function $\mathrm{erf}(\lambda'-\tau')$ around $\lambda = 0$ becomes parallel to $\lambda$ axis, being much closer to the constant function. Then, the cross point of $\mathrm{erf}(\lambda'-\tau')$ and $h$ gets highly sensitive to small changes of $p$ value, looks more and more similar to the discontinuous transition. Especially, under the $\sigma \to 0$ limit, the error function converges to the step function $\Theta(\lambda-\tau)$ that the intersection with $h$ is possible only at $\lambda = \tau$ except $p = 0.5$ at which $(1-p)/p$ completely overlaps to the step function in the region $\lambda < \tau$. Note that now the antibiotics application time is $\tau$ with no fluctuation, and thus, the transition of the optimal lag time from 0 to $\tau$ corresponds to the discontinuous transition discussed in the previous section.

#### 10.3 A sufficient and necessary condition for the discontinuous transition

We showed that if and only if the support of Normal distribution has non-zero lower bound, the optimal lag time exhibits a discontinuous transition. Indeed, exactly the same argument is applied for an arbitrary probability distribution functions which has a well-defined mean.

Suppose any distribution function $q(T)$ with its support $(T_{\rm lb}, T_{\rm ub})$ where $T_{\rm ub}$ can be either finite or infinite. Here we assume that $q(T)$ has a well-defined mean. The first-order derivative of the fitness function $\mathcal{F}_I^\delta$ respect to $\lambda$ is given as

$$
\begin{aligned}
\frac{\partial \mathcal{F}_I^\delta}{\partial \lambda} &= \begin{cases} -(1-p)+\gamma p & (\lambda < T_{\rm lb}) \\ p(1+\gamma)\Big(-\mathcal{G}_I^\delta(\lambda)+\mathcal{H}_I^\delta(\gamma,p)\Big) & (T_{\rm lb} \leq \lambda < T_{\rm ub}) \\ -1 & (T_{\rm ub} \leq \lambda), \end{cases} \\
\mathcal{G}_I^\delta &= Q(\lambda) \\
\mathcal{H}_I^\delta &= \frac{1}{1+\gamma}\Big(Q(T_{\rm lb})+\gamma Q(T_{\rm ub})-\frac{1-p}{p}\Big)
\end{aligned}
$$

where $Q$ is an arbitrary chosen primitive function of the distribution function of $q$. Again, the optimal lag time is zero as long as $-(1-p)+\gamma p < 0$ holds, and otherwise, it transits discontinuously to a non-zero value determined by the second case of above equation.

To trigger the transition of the optimal lag time from zero to non-zero, $\mathcal{H}_I^\delta$ must approach to $\mathcal{G}_I^\delta$ from below because for any $\gamma, p$ values, there are values of $\lambda$ such that $\mathcal{G}_I^\delta \geq \mathcal{H}_I^\delta$ holds. Since $q(T) \geq 0$ for $^\forall T \in \mathrm{supp}(q)$ and $Q(T)$ is a primitive function of $q(T)$, $Q(T)$ takes its minimum at $T = T_{\rm lb}$, and thus, the first intersection of $\mathcal{G}_I^\delta$ and $\mathcal{H}_I^\delta$ is formed at $\lambda = T_{\rm lb}$, and moves continuously with changes of the parameter values. Thus, the condition is sufficient.

Especially, in the case of $T_{\rm lb} = 0$, the transition becomes continuous. By taking the contraposition of the argument, the condition $T_{\rm lb} > 0$ is now shown as necessary.
